# A new model of spinal cord injury by cryoapplication: Morphodynamics of histological changes of the spinal cord lesion

**DOI:** 10.1101/2020.09.09.289025

**Authors:** George B. Telegin, Alexey N. Minakov, Aleksandr S. Chernov, Vitaly A. Kazakov, Elena A. Кalabina, Alexey A. Belogurov, Nikolay A. Konovalov, Aleksandr G Gabibov

## Abstract

Up to 500,000 people worldwide suffer from spinal cord injuries (SCI) annually, according to the WHO. Animal models are essential for searching novel methodological guidelines and therapeutic agents for SCI treatment. We developed an original model of posttraumatic spinal cord glial scar in rats using cryoapplication. The method is based on cryodestruction of spinal cord tissue with liquid nitrogen. Thirty six male SD linear rats of SPF category were included in this experimental study. A T13 unilateral hemilaminectomy was performed with an operating microscope, as it was extremely important not to penetrate the dura mater, and liquid nitrogen was applied into the bone defect for one minute. The animals were euthanized at various intervals ranging from 1 to 60 days after inducing cryogenic trauma, their Th12-L1 vertebrae were removed “en bloc” and the segment of the spinal cord exposed to the cryoapplicator was carefully separated for histological examination. The study results demonstrated that cryoapplication of liquid nitrogen, provoking a local temperature of approximately minus 20°C, produced a highly standardized transmural defect which extended throughout the dorsoventral arrangement of the spinal cord and had an “hour-glass” shape. During the entire study period (1-60 post-injury days), the glial scarring process and the spinal cord defect were located within the surgically approached vertebral space (Th13). Unlike other available experimental models of SCI (compression, contusion, chemical, etc.), the present option is characterized by a minimal invasiveness (the hemilaminectomy is less than 1 mm wide), high precision and consistency. Also, there was a low interanimal variability in histological lesions and dimensions of the produced defect. The original design of cryoapplicator used in the study played a major role in achieving these results. The original technique of high-precision cryoapplication for inducing consistent morphodynamic glial scarring could facilitate a better understanding of the self-recovery processes of injured spinal cord and would be helpful for proposing new platforms for the development of therapeutic strategies.

## Introduction

Spinal cord injury (SCI) is one of the main causes of disability associated with the inevitable formation of a glial scar in the posttraumatic period, which impedes the regenerative axonal growth through the site of the lesion [1–3]. An extensive research of potential therapeutic solutions has not provided promising results so far [4–5], though some of the recently proposed therapeutic protocols, such as epidural stimulation, seem to be more encouraging [6–8]. Therefore, it appears evident that reliable and easily reproducible animal models are needed for testing potential treatments.

Rodent models are typically used in the experiments with SCI induction to assess compression- or contusion-simulating impact [9–11]. Though these SCI models reproduce a realistic clinical course of the spinal cord injury in humans, they have multiple drawbacks, in particular, the impossibility to induce a «standardized» defect. In this context, the glial scarring process in the experimental SCI simulation is accompanied by a poor reproducibility. This limitation poses a serious challenge for proposing therapeutic agents (treatments) based on the experimental data.

Since post-traumatic glial scars either fully prevent or significantly affect the growth of axons through the defect site after SCI [12], it seems to be important and promising to find novel methods of SCI simulation that would ensure a reliable reproducibility of a standardized controlled glial scar. Our research group has recently described a new technique of SCI induction in experimental rats which is based on the local cryodestruction to produce a standardized glial scar [13]. The goal of the present study is to review the time course of histological changes and the formation of a glial scar using the proposed experimental SCI model.

## Results

### Acute period after SCI

The macro- and microscopic patterns of the damaged site were inspected and compared with the contralateral non-injured area. Macroscopically the injured area differed significantly from the non-damaged side of the cord (Fig 1) and was always located at the level of the surgical approach (Fig 2).

**Fig 1.**
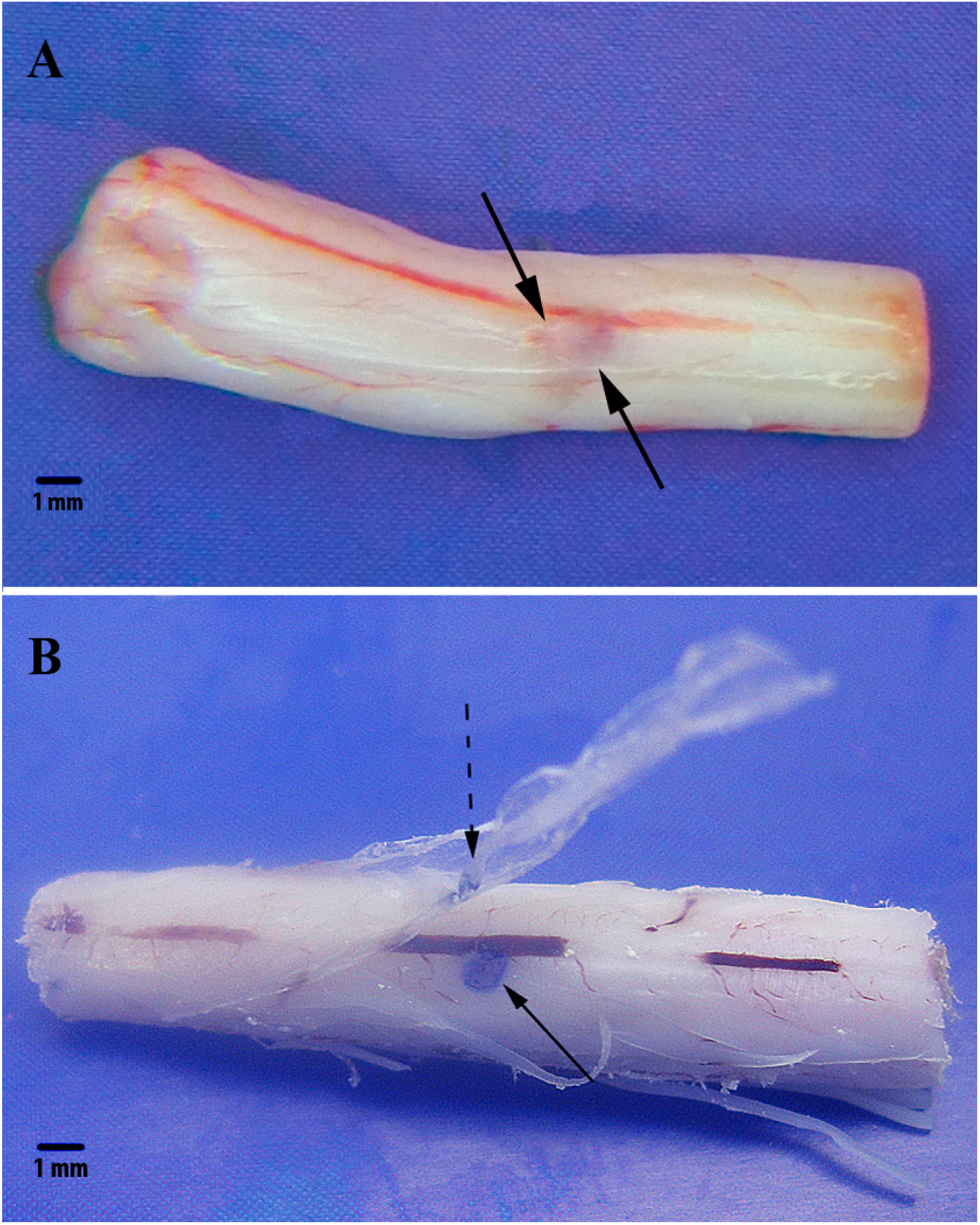
Isolated sample of the spinal cord of a rat 1 day after cryoapplication. A – macroscopical appearance of the lesion in the area of cryoapplication. Scale bar, 1 mm. B – Methylene blue staining gives a clearer view of the lesioned area (solid arrow) as well of the adjacent dura mater (dotted arrow).

**Fig 2.**
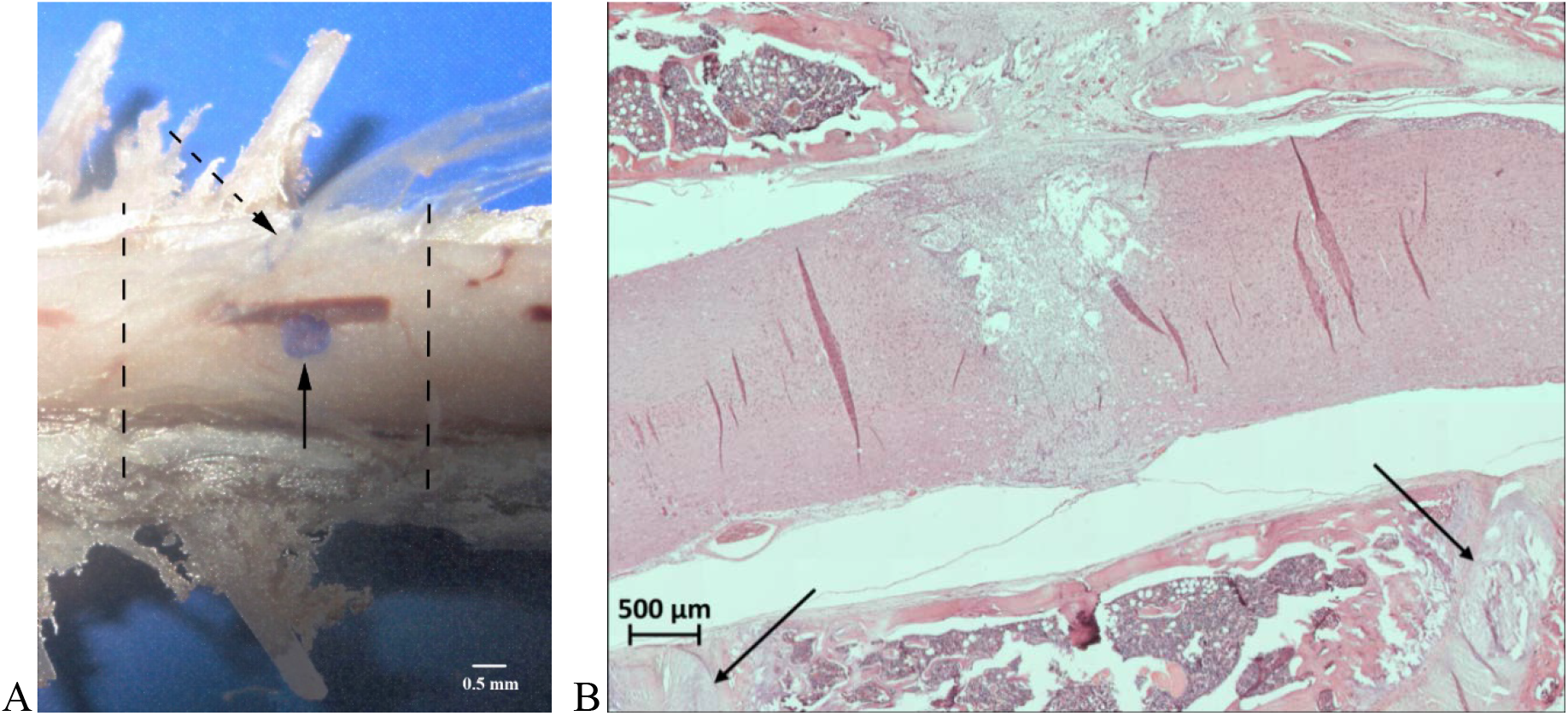
The macro- and microscopic patterns of the damaged site of spinal cord. A - sample of the spinal cord together with bone tissue. Defect of the spinal cord (solid arrow); cryoapplication zone on the dura mater (dotted arrow); margins of Th13 vertebra (dotted lines). Scale bar, 0,5 mm. B - sagittal section of Th13 vertebra after cryoapplication. The lesioned spinal cord within the projection of the approached vertebra. The upper and lower margins of T13 are shown by arrows. Scale bar, 500 μm.

It is worth noting that the injured area of the spinal cord was strictly limited by the projection of the site of the laminectomy -Th13 vertebra - (Fig 2).

During the first day after the cryoinjury, some typical signs of acute ischemic lesion were found in the affected area: tissue debris; massive microhemorrhages with a pronounced imbibition of the spinal cord tissues by fresh erythrocytes; few segmented neutrophils entering the site of necrosis; neutrophil margination and diapedesis in the vessels close to the necrotic zone.

### Subacute period after SCI

In the early subacute stage (days 3-5 after the trauma), the cryodestruction site was characterized by well-defined boundaries separating it from the surrounding healthy tissue of the spinal cord. At day 3, noticeable hemorrhagic signs were still present in the cryodestruction site, and clusters of segmented neutrophils were spread over the central and peripheral parts of the defect.

At this stage the area of necrosis was characterized by hypocellularity. In addition, a pronounced vascular response was noticed at the interface between the damaged and the intact spinal cord tissues. Individual macrophages (siderophages) were visualized at the defect margins on the side of the intact tissue, and erythophagocytosis was also observed. In general, at this stage the activation of macrophages was relatively low.

Beginning from the fifth day following the injury, the spinal cord defect acquired its final geometric pattern of an «hour-glass» shape, expanding from the dorsal side and narrowing from the ventral side. The narrowed portion of this “hour-glass” corresponds to the anterior third of the spinal cord gray matter.

The cryodestruction site was characterized by the so-called «inhibition of leucocyte infiltration»: as a rule, very few, if any, segmented leucocytes were present in the lesioned area (Fig 3A). Some segmented neutrophiles undergoing degradation were also detected.

**Fig 3.**
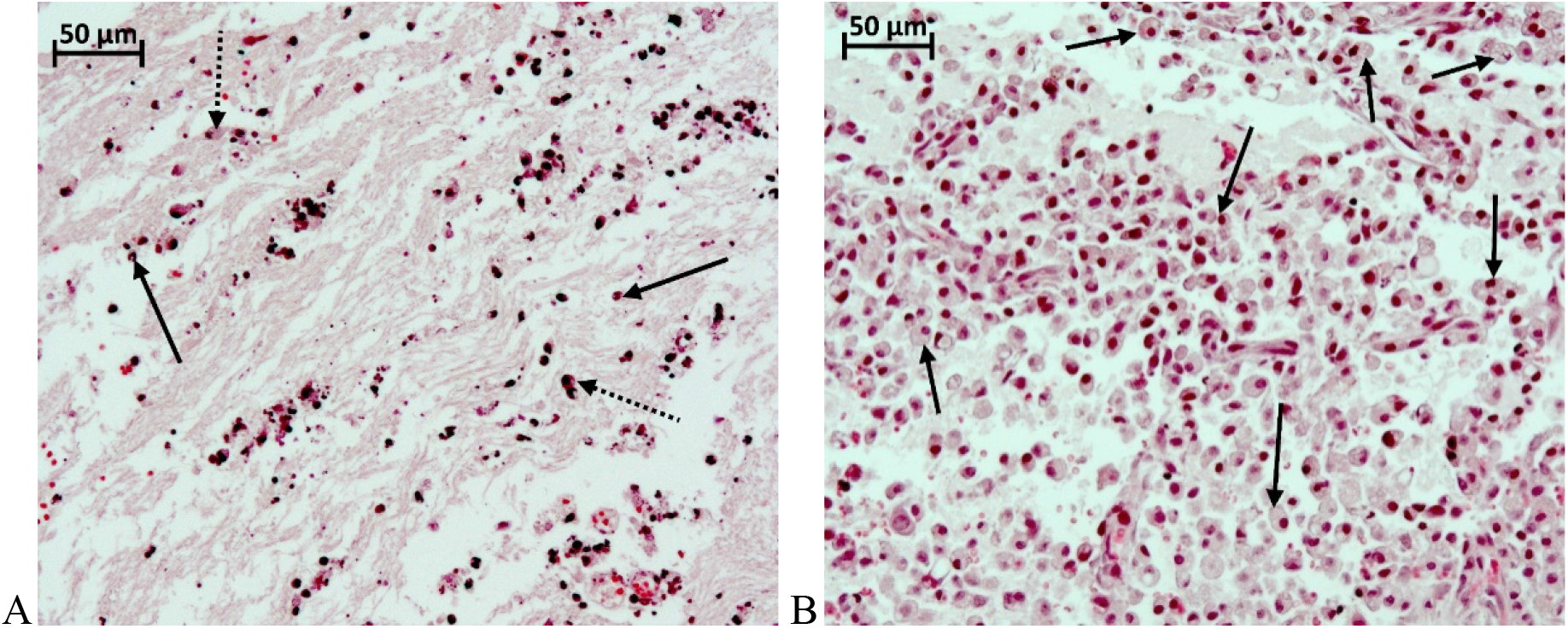
The microscopic photos of fragment spinal cord at day 5 after the cryoinjury. A - Fragment of the central portion of the necrotic area. The «inhibition of leucocyte infiltration» with individual segmented leucocytes (solid arrows) versus few macrophages (dotted arrows). Small fragments of the decomposed segmented leucocytes are also visible in the necrotic zone. B - Fragment of the peripheral part of the necrotic zone: multiple macrophages transforming into «grainy spheres» (solid arrows). H&E staining. Magnification 200x.

At the same time, the periphery of the lesion area appears to be largely infiltrated by activated macrophages. They are beginning to transform into typically looking lipophage-like «grainy spheres» due to their lipid-rich cytoplasm composition (Fig 3B). During the above time span a peak vascular response was recorded in the peripheral zone of the defect. In addition, some initial manifestations of neoangiogenesis with an active role played by fibroblasts were evident at the peripheral portion of the defect. Some signs of the proliferation of endothelial cells were also noticed. Macrophages - transformed into siderophages - were involved in the active removal of late-stage hemorrhagic components.

At day 7 after the cryoinjury, macrophages were distributed evenly throughout the induced defect of the spinal cord (Fig 4A). On this day, the number of macrophages per unit in the area of necrosis reached its maximum value: 68 cells per field of vision of 37500 μm^2^ (with 400x magnification). Manifestations of erythrophagocytosis were present both in the central and peripheral parts of the necrotic zone.

**Fig 4.**
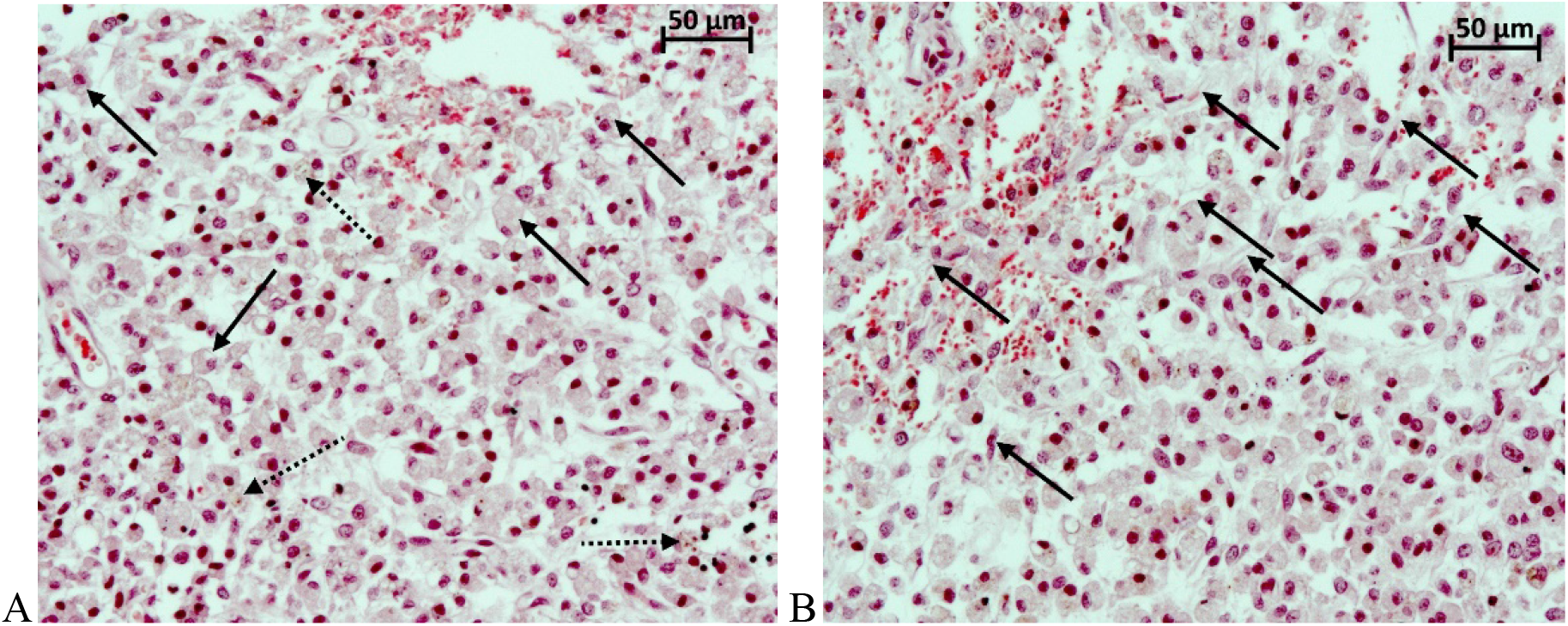
The microscopic photos of fragment spinal cord at day 7 after the cryoinjury. A - Fragment of the central part of the necrotic site: multiple macrophages transforming into «grainy spheres» and filling up the entire defect (solid arrows); multiple siderophages (dotted arrows) reabsorb the late-stage hemorrhagic component in the necrotic site; on-going erythrophagocytosis process. B - Fragment of the peripheral part of the necrotic site in the rat spinal cord interfacing with the intact tissue: multiple thin-walled blood vessels with active proliferation of the endothelium (solid arrows). H&E staining, magnification 200x.

In the central and peripheral portions of the defect, the erythrocytes were scarce and their shape looked abnormal. A high activity of neoangiogenesis accompanied by active proliferation of endothelial cells was demonstrated at the margins of the necrotic area (Fig 4B). The average specific volume of vessels in the area of defect rose to 0.0359 mm^3^/mm^3^.

In the late subacute stage (days 10-14), there was a gradual decrease in the number of macrophages in the defect. In parallel, glial cells were emerging, the process of neoangiogenesis was progressing and the proportion of collagen fibers in the scar tissue was growing. As a rule, macrophages formed clusters suggesting that the phases of gliomesodermal scarring varied in different sections of the lesioned area. The newly formed vessels were found in the central part of the defect and their specific volume increased considerably, from 0.0359 to 0.0504 mm^3^/ mm^3^. Histological structure of the vascular walls was gradually returning to its regular pattern.

At day 14 after the cryoinjury the macrophage percentage of the total cell population was still decreasing in the area of defect (to 43 cells per field of view of 37500 μm^2^) (Fig 5A); the newly emerged multiple clusters of glial cells were spread unevenly throughout the area of defect (Fig 5B). The scar tissue was characterized by regular vascularization pattern and the blood vessels acquired a common histological organization.

**Fig 5.**
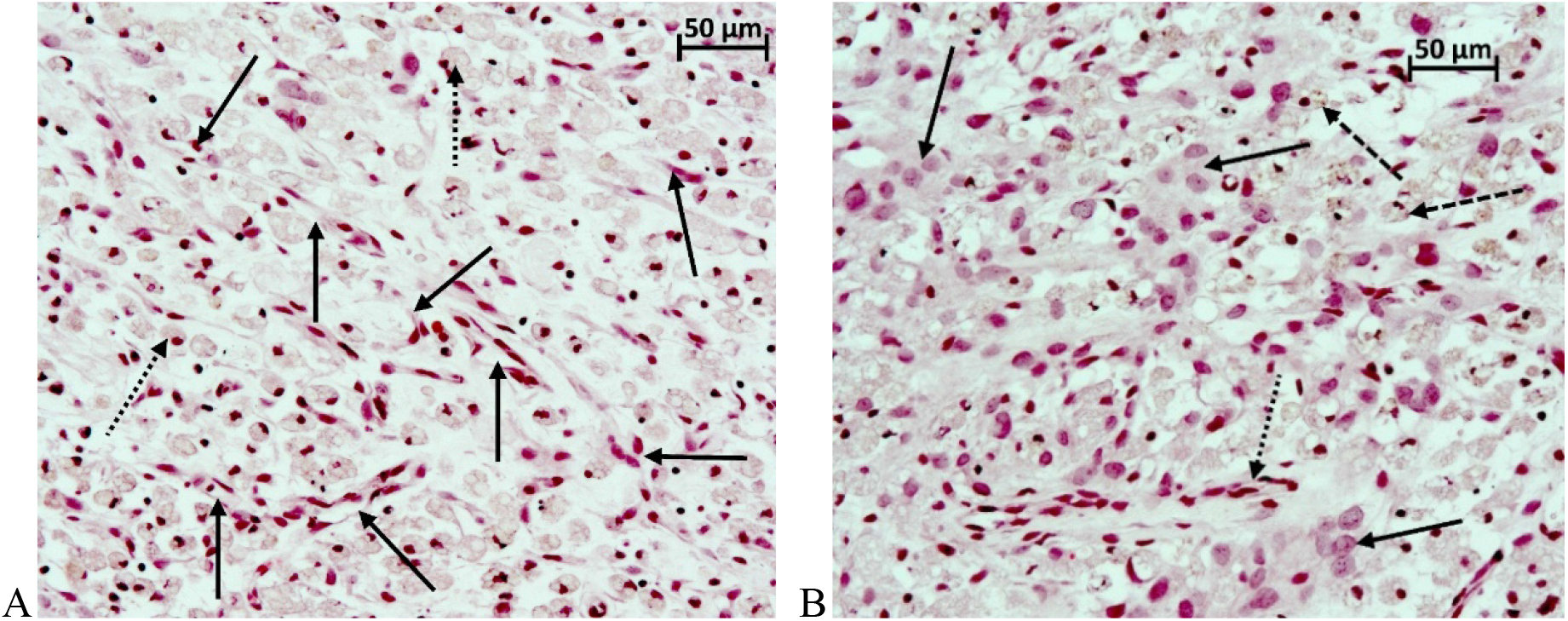
The microscopic photos of fragment spinal cord at day 14 after the cryoinjury. A - Fragment of the central part of the defect: further reduction of the number of macrophages in the overall population of cells in the lesion (dotted arrows). Multiple newly formed blood vessels (solid arrows). B - Fragment of the peripheral part of the defect: further decline of macrophages in the overall population of cells in the defect area (dashed arrows), the emergence of large glial cells (solid arrows). Newly formed blood vessels (dotted arrow). H&E staining, magnification 200x.

By day 14, the average specific volume of blood vessels in the area of defect increased significantly and reached 0.0725 mm^3^/mm^3^. At day 14 the fibrous tissue component made a very low contribution to building the structure of gliomesodermal scar. At the same time, some artificial hollow spaces were found in the histological sections. The artificial hollow spaces developed as a result of the «fragility» and looseness of the defect site.

At day 21, the number of macrophages was still high and they were distributed evenly throughout the formed defect in the spinal cord (at the average 43 cells per field of view of 37500 μm^2^). However, their morphology and size varied within the same histological sections. Clusters of glial cells were visible in the peripheral zone of the defect. The size of fibrous tissue components in the entire structure of the gliomesodermal scar was noticeably increasing on day 14, while the artificial hollow spaces in the sections were still present. It is worth noting that already at this time span a trend to the development of cystic cavities in the site of cryoinjury was found in one of the experimental animals. Multiple newly formed blood vessels with an ordinary histological structure were visible all over the area. The average specific volume of blood vessels in the site of defect reached 0.0807 mm3/mm^3^.

### Chronic period after SCI

By day 30 of the follow-up period, multiple macrophages were still present and distributed evenly throughout the site of defect of the spinal cord (averaging 46 cells per field of view of 37500 μm^2^), whilst their morphology and size varied within the same histological sections (Fig 6A). It should be highlighted that at this time interval a trend to the development of cystic cavities in the site of cryoinjury was already traced in one of the experimental animals (Fig 6B).

**Fig 6.**
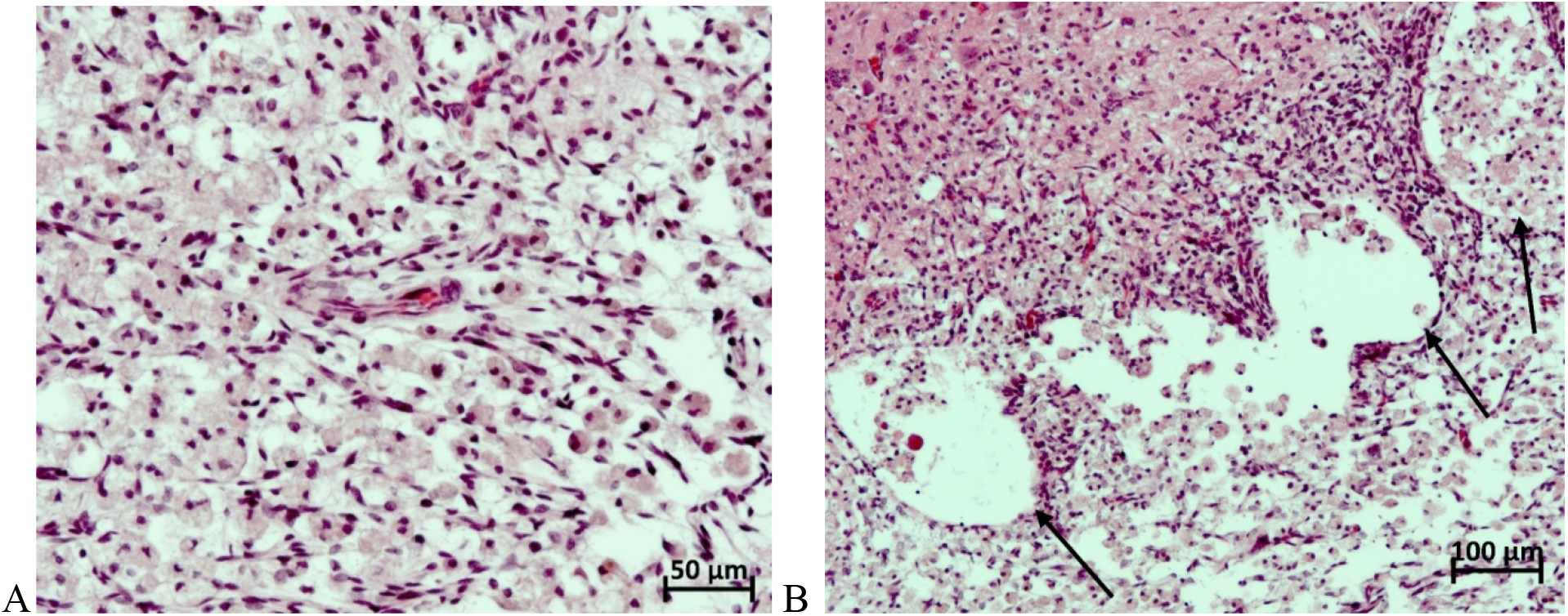
The microscopic photos of fragment spinal cord at day 30 after the cryoinjury. A - Fragment of the central part of the defect: plenty of macrophages distributed throughout the defect area on the background of multiple newly formed blood vessels. B - Fragment of the peripheral part of the necrotic zone in the rat spinal cord, interfacing with the intact tissue: a trend toward the formation of multiple small cystic cavities at the border between the defect and the intact spinal cord, filled up mainly with macrophages. H&E staining, magnification 200x.

The proportion of fibrous tissue component was increasing gradually in the structure of gliomesodermal scar, while the artificial hollow spaces were markedly less present in the histological sections than at days 14 and 21 after the cryoinjury. Multiple newly formed blood vessels with a normal histological structure were spread all over the examined areas. The average specific volume of blood vessels in the site of defect reached 0.0878 mm^3^/mm^3^.

Two months after the cryoinjury, in all animals the spinal cord defect contained large cystic cavities filled up with an amorphous substance which could reach the size of 0.8 mm^2^. The share of fibrous tissue component in the entire structure of the gliomesodermal scar was rather low, the unevenly distributed fibers were undergoing different stages of maturity.

Cellularity in the area of defect was also non-homogeneous with a moderate macrophage-predominant pattern. Few glial cells and fibers are located at the defect margins (Fig 7A). By the end of the second month, multiple macrophages were still found but in a significantly reduced quantity (from 0 to 40 cells per field of view of 37500 μm ^2^, averaging 21 cells) (Fig 7B). In the area of the spinal cord trauma the vascular system was well-developed and blood vessels acquired a normal histological structure.

**Fig 7.**
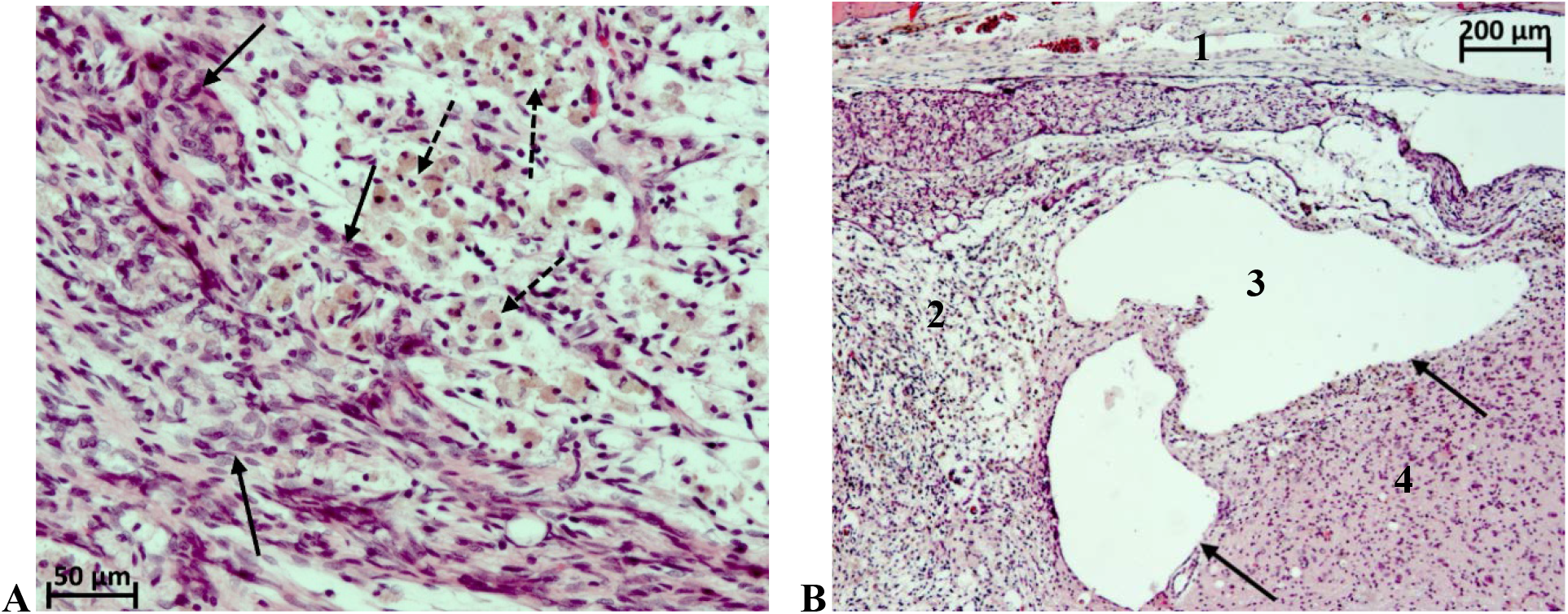
The microscopic photos of fragment spinal cord at day 60 after the cryoinjury. A - Fragment of the peripheral part of the defect: multiple macrophages (dotted arrows) on the background of glial cells and glial fibers. B - Fragment of the peripheral part of the necrotic zone, interfacing with the intact tissue: large cystic cavities at the border between the defect and the intact spinal cord (1 - Approach area, 2 - Defect of the spinal cord, 3 - Cystic cavities, 4 - Intact spinal cord). H&E staining, magnification 200x.

Figure 8 summarizes the timeline of changes in macrophage number and specific volume of blood vessels in the lesioned area of the assessed experimental material.

**Fig 8.**
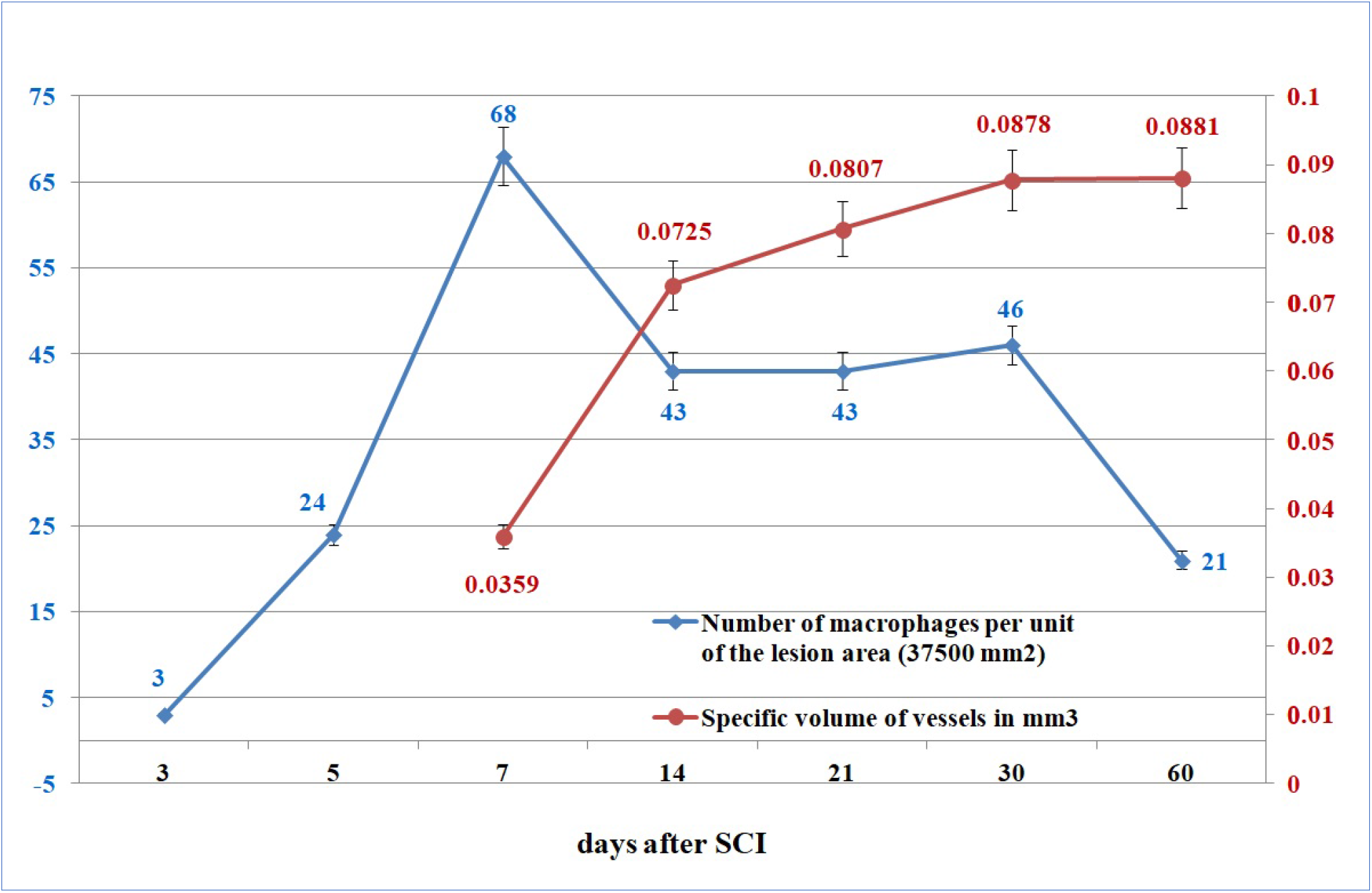
Changes in the numbers of macrophages and the volume of blood vessels in the cryolesion area in experimental rats.

Impact of the cryoapplication on locomotor activity of rats was tested in the “open field” according to 21-score BBB Locomotor Rating Scale. Animals from the experimental group demonstrated stable monoplegia with abnormalities of locomotor functions at a mean 2.7 BBB score for 1 to 60 days. In the control group of animals where only surgical approach to the spinal cord was performed (without exposure to low temperatures), the full recovery of motor activity was reported much earlier – five days after the surgery (fig 9).

**Fig 9.**
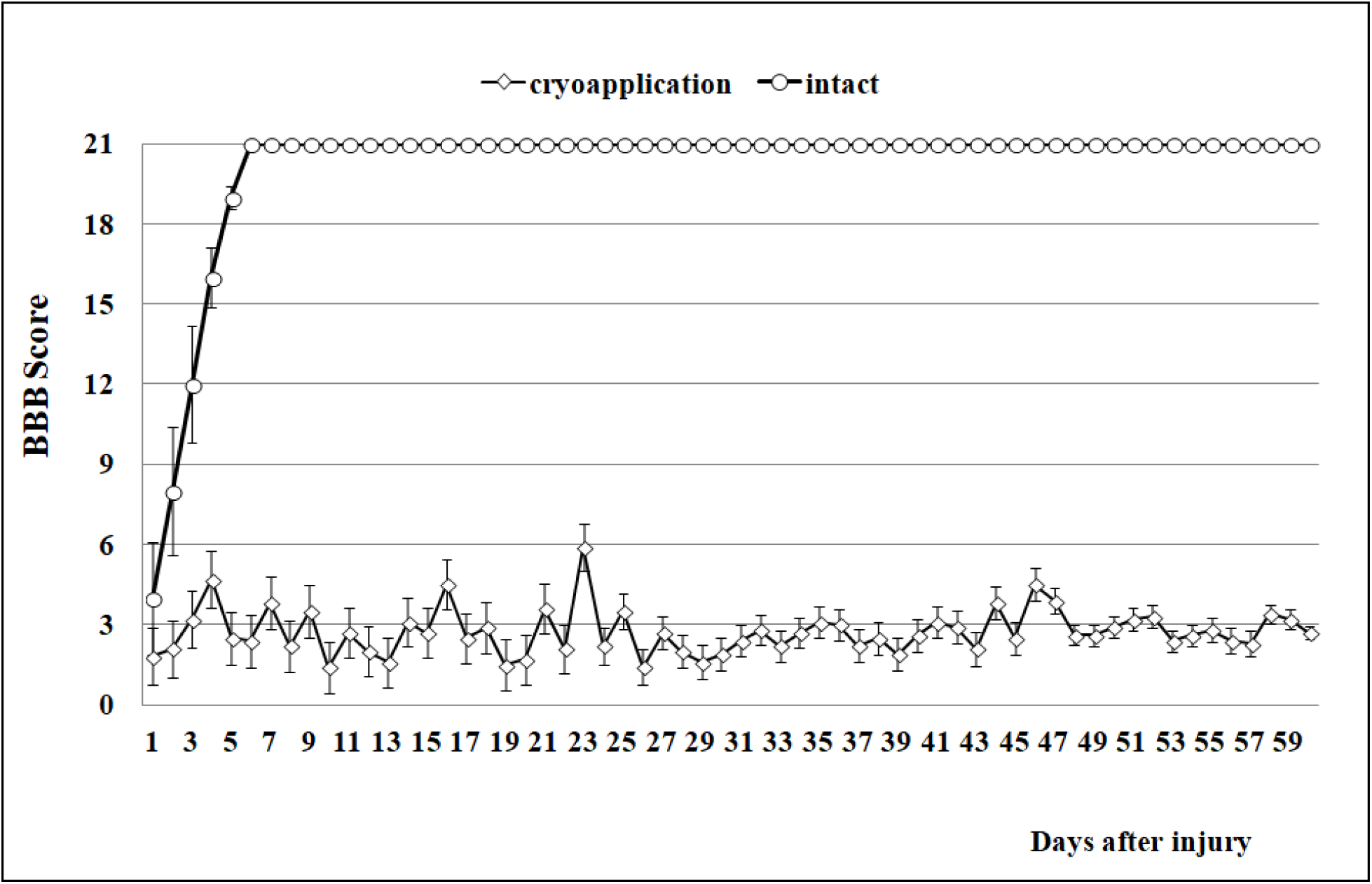
Mean BBB (Y axis) score of the rats over the course of 1 month follow-up (X axis) demonstrating minimal recovery typical for severe SCI.

## Discussion

This histological study provided clear data on the time course of glial scar formation following a standardized cryoinjury of the rat spinal cord. In general, the observed scarring process showed an expected sequence of events. Histological findings demonstrated that the cryodestruction sites in serial sagittal sections were transmural, i.e. they encompassed the dorsoventral arrangement of the spinal cord along its entire length. Formation of an «hourglass-shaped» spinal cord lesion (which appeared to be highly reproducible in multiple animals) with tissue necrosis was observed in the acute postinjury period. The spinal cord defect induced by cryoapplication comprises the right dorsal, right lateral and right ventral funiculi of spinal white matter, as well as the right dorsal and ventral horns of grey matter that ensures a stable monoplegia not followed by self-recovery. It is worth noting that the commonly used methods of SCI simulation do not meet this requirement [11, 14]. Current SCI models cause significant urinary system dysfunctions in rats, which is a serious drawback [14]. It is necessary to manually empty the bladder of the animals several times a day after injury to avoid bladder rupture and infectious inflammation [15–16]. Our model had no such drawback thanks to the minimal surgical injury. After injury, the animals retained their ability to naturally empty their bladder and intestines during the entire follow-up period despite the persistent monoplegia. The ability to urinate independently and defecate is the key to life support in the chronic postoperative period; it prevents the development of distress in rats and a nonspecific injury to the spinal cord when stimulating natural movements by palpation on the walls of the intestines and bladder through the abdominal wall of the animal.

The subacute period was characterized by the macrophage-mediated reabsorption of necrotic tissue. This process was followed by local neoangiogenesis and ultimately resulted in the development of mesodermal-glial scarring tissue and the formation of cysts. For the first time, macrophages appeared at the peripheral portion of the defect on day 3 and demonstrated an exponential growth by day 7. However, their number reduced by day 14 and reached a plateau at which it stayed at least by the end of the first month of follow-up period. It should be pointed out that at day 60 multiple macrophages were still present in the defect site. These findings definitely prove that rearrangement processes occur continuously in the process of maturation of a glial scar [12, 17]. Neoangiogenesis in the peripheral portion of the defect was already noticeable by day 5. The newly formed blood vessels begin to acquire normal histological features by day 14 of follow-up period; their specific volume demonstrates an exponential progression and reaches plateau by day 30. An adequate nutrition of the forming scar is crucial at the early stages. But it is equally important that the newly formed blood vessels keep functioning at the late stages of maturation of a glial scar [12, 18]. In our opinion, quantitative and especially qualitative indicators of the pass-through capacity of newly formed blood vessels may be considered as key predictors of the effectiveness of repair processes in nerve tissue. Thus, our future research will be dedicated to neoangiogenesis and characterization of vascular networks in the scar tissue.

Thus, the present model can be used as an efficient and reliable experimental simulation of glial scarring in the posttraumatic period. It is also noteworthy that the spinal cord lesions were usually characterized by high reproducibility and significantly decreased interanimal variability. According to BBB score, most of the rats with simulated post-traumatic scar of the spinal cord developed monoplegia at the affected side persisting for 60 days. In this context, the proposed model looks even more interesting and can be used for a reliable assessment of the process of post-SCI glial scar formation. In addition, it could be recommended for testing potential therapeutic solutions.

However, it is obvious that the proposed model does not represent the scenario of pathophysiologic events of spinal trauma that typically occur in the clinical setting. For the reproduction of clinical cases the researchers usually use other well-known experimental SCI models that cause a blunt trauma.

But we would like to re-emphasize that this model is focused on the issue of posttraumatic glial scarring, one of the most critical consequences of spinal trauma, for the management of which no successful therapeutic solutions have been found so far [19–21]. The model proposed in this study offers a reliable and standardized simulation of the spinal cord lesion and highlights the time course of posttraumatic changes. Thus, it can serve as an optimal platform for studying the efficacy of potential therapeutic solutions targeted at preventing the formation of glial scar in the posttraumatic period.

## Conclusion

The described model of standardized post-traumatic spinal cord glial scarring demonstrated that there was a specific course of histological changes. It can be used as a reliable experimental model for testing therapeutic solutions that prevent posttraumatic glial scarring.

## Methods

### Laboratory animals

Male SD rats of SPF-category (n=36) weighing 320–360 g were used in the experiment to ensure an adequate, convenient visualization and identification of all anatomical structures while performing the surgical procedure. The animals were kept under standard housing conditions at the Animal Breeding Facility of the Branch of the Institute of Bioorganic Chemistry (unique scientific unit «Bio-model» IBCh RAS). All animals manipulations were approved by the Institutional Animal Care and Use Committee of the BIBCh (Protocol No. 718/19 of 01/10/19).

### Preoperative preparation and anesthetic support

The animals were placed in cages with clean bedding and water 24 h before the surgery. The surgical procedure was performed under general anesthesia with Aerrane (Baxter Healthcare Corp., USA) on a temperature-controlled operating table (+38°C). A single intramuscular injection of Baytril (enrofloxacin, 25 mg/mL) was administered at the dosage of 10 mg/kg. Premedication was not used. Blood pressure and vital parameters were continually monitored and the surgical procedure was carried out under strictly controlled aseptic conditions.

### Surgical approach and cryoapplication

The surgical technique was described in detail in previous papers [22–23]. Briefly, unilateral laminectomy of Th13 vertebra was performed with a dental burr and cryoinjury was induced using an original device. The spinal cords of the experimental animals were cooled by applying a cryoconductor through the dura mater. In the area of contact with the biological substrate, the copper conductor was 0.8 mm in diameter. The distance from the cold source - liquid nitrogen - was 9 cm, and the cryoapplication procedure lasted for 1 min. In the contact area the local temperature reached minus 20°C.

### Postoperative monitoring

Animals were euthanized at various time intervals following the cryoinjury: in the acute (first 24 hours after the injury) subacute (days 3, 5, 7, 10, 14), and chronic period (days 21, 30, 60) (4 animals at each time interval). The present SCI protocol and clinical monitoring ensured a 100% survival rate of the experimental animals. In particular, no infectious complications were reported.

### Assessment of locomotor activity

Locomotor activity of rats was tested prior to surgery, and every day after injury for 60 days. Two independent observers who were blinded to the treatment methods and groups performed Basso, Beattie and Bresnahan (BBB) scoring in an open field test [24]. These tests included 4 rats per group and were conducted to evaluate the restoration of hindlimb locomotor function after SCI. For the BBB scoring, the rats were individually placed in an open area with a non-slippery surface and allowed to move freely for 5 min. Two observers evaluated locomotion during open-field walking and scored the hindlimb performance according to the BBB scale in which scores range from 0 (no movement) to 21 (normal movement).

### Histological methods

Samples of the rat injured spinal cord were removed “en bloc” together with three vertebrae (the vertebra of the surgical approach – Th13, and two adjacent vertebrae – Th12 and L1). They were fixed in 10% neutral buffered formalin solution for 2-5 days, rinsed in running tap water and processed for decalcification in Trilon B at a room temperature for 12-16 days.

As soon as a satisfactory decalcification was achieved, the specimens were cut and cleaned of soft tissues, and the biomaterial was oriented for further microtomy in the sagittal plane. The cut specimens were rinsed in running tap water, dehydrated in an ascending alcohol series and embedded in paraffin. Microtomy of the specimens was performed in the sagittal plane, in series with a sectioning step size of 200 μm.

Hematoxylin and eosin (H&E) as well Heidenhain's azocarmine were used for staining serial 4-5-μm-thick paraffin sections. A series of sections in the sagittal plane was prepared for morphological measurements, and the section with the largest defect area was selected. The topography of affected structures of the spinal cord was assessed according to «A high-resolution anatomical rat atlas» [25].

In order to improve visualization of the spinal cord defect and to assist with preparing histological sections, a methylene blue 0.1% solution (HiMedia Laboratories, India) was used. After the animals were euthanized and the spinal cord within the bone fragment was extracted, 100 μl of dye were added to the laminectomy window. As a result, an intense blue staining of the defect area was achieved.

Then, the sections were examined by ordinary light microscopy using Zeiss Axio Scope A1 microscope (Carl Zeiss, Germany). Microphotographs of the histological sections were made with Axiocam 305 color high-speed camera (Carl Zeiss, Germany), and morphometric measurements were processed with ZEN 2.6 lite software (Carl Zeiss, Germany). The obtained data relaying to the number macrophages and the volume of blood vessels were processed using a statistical software package SigmaPlot statistic (v. 13.0).

